# The first multi-tissue diel genome-scale metabolic model of a woody plant highlights suberin biosynthesis pathways in *Quercus suber*

**DOI:** 10.1101/2021.03.09.434537

**Authors:** Emanuel Cunha, Miguel Silva, Ines Chaves, Huseyin Demirci, Davide Rafael Lagoa, Diogo Lima, Miguel Rocha, Isabel Rocha, Oscar Dias

**Affiliations:** Centre of Biological Engineering, Universidade do Minho, 4710-057 Braga, Portugal; LABBELS – Associate Laboratory, Braga, Guimarães, Portugal; Instituto de Tecnologia Química e Biológica António Xavier, Universidade Nova de Lisboa, Avenida da República, Quinta do Marquês, 2780-157 Oeiras, Portugal; iBET, Instituto de Biologia Experimental e Tecnológica, Apartado 12, 2781-901 Oeiras, Portugal; SnT/University of Luxembourg, Luxembourg

**Keywords:** Cork Biosynthesis, *Quercus suber* (Cork Oak), Genome-scale Metabolic Model, Multi-tissue diel cycle model, Secondary Metabolism, Suberin

## Abstract

In the last decade, genome-scale metabolic models have been increasingly used to study plant metabolic behavior at the tissue and multi-tissue level under different environmental conditions. *Quercus suber*, also known as the cork oak tree, is one of the most important forest communities of the Mediterranean/Iberian region. In this work, we present the genome-scale metabolic model of the *Q. suber* (iEC7871), the first of a woody plant. The metabolic model comprises 7871 genes, 6231 reactions, and 6481 metabolites across eight compartments. Transcriptomics data was integrated into the model to obtain tissue-specific models for the leaf, inner bark, and phellogen, with specific biomass compositions. The tissue-specific models were merged into a diel multi-tissue metabolic model to predict interactions among the three tissues at the light and dark phases. The metabolic models were also used to analyze the pathways associated with the synthesis of suberin monomers. Nevertheless, the models developed in this work can provide insights into other aspects of the metabolism of *Q. suber*, such as its secondary metabolism and cork formation.

## 1. Introduction

The cork oak, *Quercus suber* L., is a characteristic tree of the Mediterranean/Iberian landscape ecosystem. The tree forms a thick bark of cork (phellem) containing high levels of aliphatic suberin, aromatic suberin, extractives (waxes and tannins), and polysaccharides (Graça, 2015). Cork properties are so unique that this material is used in diverse applications from the common bottle-stopers to spacecrafts built by NASA and European Space Agency (ESA), providing insulation solutions to deal with extreme conditions. The aliphatic suberin component has particular interest since it provides waterproof, light, elastic, and fire-retardant properties to cork (Vaz *et al*., 2011; Pereira, 2015). Only the outer bark is separated from the trunk during the harvest, which enables the regeneration and allows using the tree as a renewable biological resource. Cork extraction can continue in the same tree for more than a century every 9 years. A cork oak tree with 234 years old, debarked 20 times, is now a living icon being elected European tree of the year 2018 (https://www.treeoftheyear.org/Previous-Years/2018).

Phellem is produced by the phellogen (cork cambium), involving the proliferation of phellogen derivates cells which undergo to differentiate into cork cells through cell expansion, deposition of suberin and waxes, and an irreversible senescence program ending with cell death. The first debarking occurs when the tree has 18-25 years old. This first harvested cork, known as “virgin” cork, produced from the original phellogen, yields poor-quality cork. The original phellogen is then replaced by a traumatic phellogen that proliferates to originate a new cork layer. From the third debarking onwards, a higher quality cork with high economic value is obtained - reproduction cork (or “amadia”) (Graça & Pereira, 2004). Cork growth and quality are dependent on the genome of each tree but it is also strongly dependent on the environment, such as water availability, temperature, pests, and diseases.

The cork oak tree represents one of the most relevant broadleaved forest resources in the Mediterranean basin (Portugal, Spain, Algeria, Morocco, France, Italy, and Tunisia), having a substantial socio-economic and ecological impact on these countries. This tree species can generally live over 200 years, which implies a capacity to bear with biotic stresses like bacteria, fungi and insects, as well as abiotic stresses like drought, floods and fires (Acácio *et al*., 2007; Kim *et al*., 2017).

Due to its economic interest in Portugal, a national consortium was created to first sequence the transcriptome (Pereira-Leal *et al*., 2014) and recently to full sequence the genome of *Quercus suber* (Ramos *et al*., 2018). The genome sequence’s availability allows the application of systems biology tools to study this species’ metabolic behavior.

Genome-scale metabolic (GSM) models aim at depicting the whole metabolic network of an organism. Such models have been widely used for metabolic engineering purposes, mainly with prokaryotes and yeasts. Among other applications, GSM models can be used to analyze an organism’s metabolic behavior in different environmental and genetic conditions, including the effect of gene knock-outs and over/under-expressions (Dias & Rocha, 2015). Curated models have proven to accurately predict complex organisms’ metabolic behavior in diverse areas of knowledge, from biotechnological or environmental to medical applications (Fong *et al*., 2005; Bordbar *et al*., 2011; Agren *et al*., 2014; Jerby-Arnon *et al*., 2014; Zhang & Hua, 2016). Genome-scale modelling in plants is much more challenging than in prokaryotes. The struggle starts in the annotation of the complex genomic content, in which the function of a significant number of genes is still unknown. Also, the subcellular compartmentalization, tissue differentiation, and interactions between tissues are complex in plants. Nevertheless, in the last decade, the number of published GSM models of plants has increased considerably. Organisms, like *Arabidopsis thaliana* (Dal’Molin *et al*., 2010; Cheung *et al*., 2013), *Zea mays* (maize) (Saha *et al*., 2011; Seaver *et al*., 2015), *Oryza sativa* (rice) (Poolman *et al*., 2013; Lakshmanan *et al*., 2013) have more than one model available. Considering the importance of plants in terms of nutrition, biofuels, and their capability to produce a variety of secondary metabolites, it is not surprising that studies for plant genome-scale model reconstruction will increase, parallel to the growth in the number of sequenced species. The utilization of user-friendly tools designed for this purpose, like *metabolic models reconstruction using genome-scale information* (*merlin)* (Dias *et al*., 2015), accelerates the reconstruction process.

This paper describes the first reconstruction of iEC7871, a Genome-scale metabolic model for the cork oak tree, and as far as we know, the first published GSM model of a tree. Besides the generic GSM, we present tissue-specific models and a multi-tissue metabolic model that can be used to study the metabolic behavior of *Q. suber*.

## 2. Results

### 2.1. Genome Annotation

A GSM model for *Q. suber* based on an up-to-date genome annotation was reconstructed in this work. A Basic Local Alignment Search Tool (BLAST) (Altschul *et al*., 1990) search against the Swiss-ProtKB (Apweiler *et al*., 2011) database allowed to identify similarity results for 47,199 out of the 59,614 genes available in the genome (Ramos *et al*., 2018). A second BLAST search against UniProtKB/TrEMBL allowed obtaining hits for 12,415 genes, while 590 had no results.

As detailed in the Materials and Methods section, *merlin*’s “automatic workflow” tool annotates genes based on the homologous gene records taxonomy. Most enzyme-encoding genes were annotated based on *A. thaliana* gene records (82%). Organisms not included in the annotation workflow represent 16% of gene annotations, and only 1% were annotated using *Quercus* genus’ gene records. A more detailed analysis of the genome annotation is available in Supplementary File 1.

### 2.2. Model Properties

The metabolic model is mass balanced and can predict growth in phototrophic and heterotrophic conditions. These conditions were defined by setting the photon and CO_2_ (phototrophic) or sucrose (heterotrophic) as the sole energy and carbon sources, respectively. Additionally, the model requires *NH*_3_ or *HNO*_3_, *H*_3_*PO*_4_, *H*_2_*SO*_4_, *Fe*^2+^, and *Mg*^2+^ to produce biomass.

The general properties of the cork oak model and other five published plant models – *A. thaliana* (Chung *et al*., 2013; Seaver *et al*., 2015), *Z. mays* (Saha *et al*., 2011; Seaver *et al*., 2015), and *Glycine max* (soybean) (Moreira *et al*., 2019) - are presented in table 1.

**Table 1.** General properties (genes, reactions, metabolites, and compartments) of the Q. suber model, and other five published plant GSM models. Only generic models were considered here.

|  | *Quercus suber* (this work) | *Arabidopsis thaliana* (Chung *et al.*, 2013) | *Arabidopsis thaliana* (Seaver *et al.*, 2015) | *Zea mays* (Saha *et al.*, 2011) | *Zea mays* (Seaver *et al.*, 2015) | *Glycine max* (Moreira *et al.*, 2019) |
| --- | --- | --- | --- | --- | --- | --- |
| **Genes** | 7871 | 1475 | 5184 | 1563 | 13,279 | 6127 |
| **Reactions** | 6231 | 1895 | 6399 | 1970 | 6458 | 2984 |
| **Metabolites** | 6481 | 1761 | 6236 | 2129 | 6250 | 2814 |
| **Compartments** | 8 | 6 | 9 | 6 | 9 | 5 |

The Cork Oak model comprises 7871 genes, 6231 reactions (enzymatic, spontaneous, and transport), 708 exchange reactions, and 6476 metabolites distributed across eight subcellular compartments – the cytoplasm, mitochondria, plastid, endoplasmic reticulum, Golgi apparatus, vacuole, peroxisome, and the extracellular space. The number of genes, reactions, and metabolites of the Cork Oak model are only comparable with the *A. thaliana* and *Z. mays* models developed by Seaver (Seaver *et al*., 2015).

The reactions and metabolites present in the cork oak model were compared with the ones present in the models developed by Chung for *A. thaliana* (Chung et al., 2013), and Saha for *Z. mays* (Saha *et al*., 2011) (Supplementary File 2), as the reactions and metabolites available in these models have the same identifiers (KEGG (Kanehisa & Goto, 2000)) as our model.

A total of 3196 unique reactions and 2883 unique metabolites were identified across the three models. The cork oak model includes 1117 reactions that are not present in the other two - Figure 1. It has 124 reactions in common only with the Maize model and 179 only with the *A. thaliana* model. The maize and *A. thaliana* models share 380 reactions, and 964 were found in the three models. The metabolites analysis shows similar behavior. This analysis is detailed in Supplementary File 2 Tables S5-S6.

**Figure 1.**
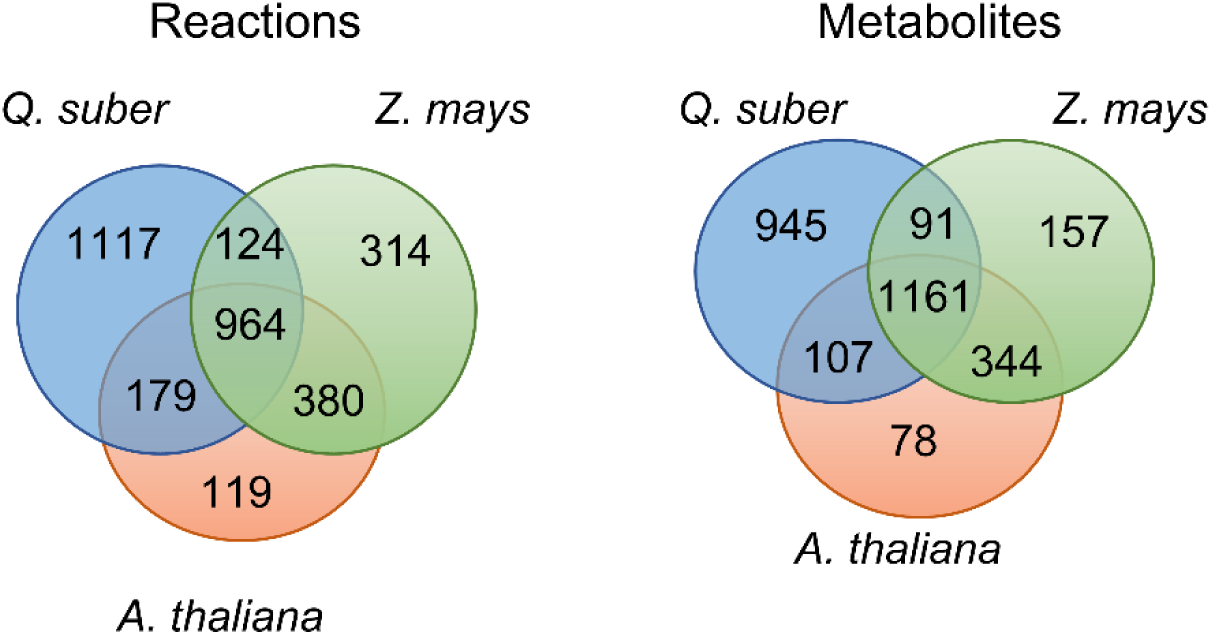
Venn diagram of reactions (left) and metabolites (right) included in the *Q. suber, A. thaliana*, and *Z. mays* models.

The reactions only available in the Cork Oak model (called unique reactions in the following) were further analyzed to assess the annotation of the genes and pathways associated with them. This analysis is detailed in Supplementary File 2 Table S7. In agreement with the remaining genome annotation, 82% of the genes encoding enzymes that catalyze these reactions were annotated based on *A. thaliana* genes’ annotations. The remaining 18% were annotated by gene records of organisms not accounted for in the annotation workflow. KEGG pathways with more unique reactions are presented in Table 2. The 34 pathways with at least 10 unique reactions can be divided into 7 main areas of the KEGG metabolism: 8 pathways representing ‘Lipid metabolism’, 7 the ‘aminoacid metabolism’ and ‘carbohydrate metabolism’, 6 the ‘Xenobiotics biodegradation metabolism’, 3 the ‘Metabolism of cofactors and vitamins’ and ‘Biosynthesis of other secondary metabolites’ and 1 the ‘Metabolism of terpenoids and polyketides’. Spontaneous reactions and reactions not associated with any pathway were not considered.

**Table 2.** Number of reactions included in the cork oak model that were not identified in the *A. thaliana* (Cheung *et al*., 2013) and *Z. mays* (Saha *et al*., 2011) models for each pathway (so-called unique reactions). Only pathways with more than 10 unique reactions were included in this table. The complete table is available in Supplementary File 2 S7. The KEGG pathways were organized according to the 7 main areas of KEGG metabolism: 1- Xenobiotics biodegradation and metabolism; 2- Lipid metabolism; 3- Metabolism of cofactors and vitamins; 4- Amino acid metabolism and Metabolism of other amino acids; 5- Metabolism of terpenoids and polyketides; 6- carbohydrate metabolism; 7- Biosynthesis of other secondary metabolites

|  | KEGG Pathway | Number of unique reactions |
| --- | --- | --- |
| 1 | Metabolism of xenobiotics by cytochrome P450 | 38 |
| 2 | Steroid hormone biosynthesis | 38 |
| 3 | Ubiquinone and other terpenoid-quinone biosynthesis | 34 |
| 4 | Tryptophan metabolism | 32 |
| 1 | Steroid biosynthesis | 24 |
| 4 | Tyrosine metabolism | 23 |
| 4 | Cysteine and methionine metabolism | 22 |
| 1 | Drug metabolism - cytochrome P450 | 21 |
| 2 | Fatty acid metabolism | 21 |
| 2 | Glycerophospholipid metabolism | 20 |
| 2 | Inositol phosphate metabolism | 19 |
| 5 | Brassinosteroid biosynthesis | 17 |
| 2 | Cutin, suberine and wax biosynthesis | 17 |
| 1 | Drug metabolism - other enzymes | 17 |
| 4,6 | Amino sugar and nucleotide sugar metabolism | 16 |
| 3 | Porphyrin and chlorophyll metabolism | 16 |
| 2 | Arachidonic acid metabolism | 16 |
| 4 | Arginine and proline metabolism | 16 |
| 3 | Nicotinate and nicotinamide metabolism | 15 |
| 4 | Cyanoamino acid metabolism | 14 |
| 6 | Butanoate metabolism | 14 |
| 7 | Betalain biosynthesis | 14 |
| 1 | Benzoate degradation | 13 |
| 6 | Fructose and mannose metabolism | 11 |
| 6 | Glyoxylate and dicarboxylate metabolism | 11 |
| 6 | Ascorbate and aldarate metabolism | 11 |
| 2 | Sphingolipid metabolism | 11 |
| 7 | Phenylpropanoid biosynthesis | 10 |
| 7 | Flavonoid biosynthesis | 10 |
| 6 | Propanoate metabolism | 10 |
| 2 | Fatty acid biosynthesis | 10 |
| 6 | Galactose metabolism | 10 |
| 1 | Chlorocyclohexane and chlorobenzene degradation | 10 |
| 4 | Lysine biosynthesis | 10 |

The unique reactions identified are associated with 122 different KEGG pathways, from which 54 had five or fewer unique reactions. “Steroid hormone biosynthesis”, “Metabolism of xenobiotics by cytochrome P450”, “Ubiquinone and other terpenoid-quinone biosynthesis”, and “Tryptophan metabolism” are the pathways with more unique reactions, having 32 to 38 reactions unavailable in the maize and *A. thaliana* models.

Several of these reactions can be related to species-specific reactions reflecting the cork oak uniqueness and differences from non-woody plants. Although the reactions are cork oak specific, they belong to pathways already present in other species. Some unique reactions are associated with pathways essential for trees and belong to the secondary metabolism responsible for wood and cork production, such as “Phenylpropanoid biosynthesis”, “Flavonoid biosynthesis”, “Fatty acid biosynthesis”, “Metabolism of xenobiotics by cytochrome P450”, and “Cutin, suberine and wax biosynthesis”.

### 2.3. Tissue-specific models

Transcriptomics data were integrated into the generic model using troppo (Ferreira *et al*., 2020) to obtain tissue-specific models. Leaf, Inner bark, Reproduction or traumatic Phellogen, and Virgin Phellogen were selected because of their influence on tree growth and cork production.

Different biomass formulations for each tissue (Figure 2) were determined, according to experimental data available in the literature and other plant GSM models. The detailed biomass composition for each tissue is available in Supplementary File 3.

**Figure 2.**
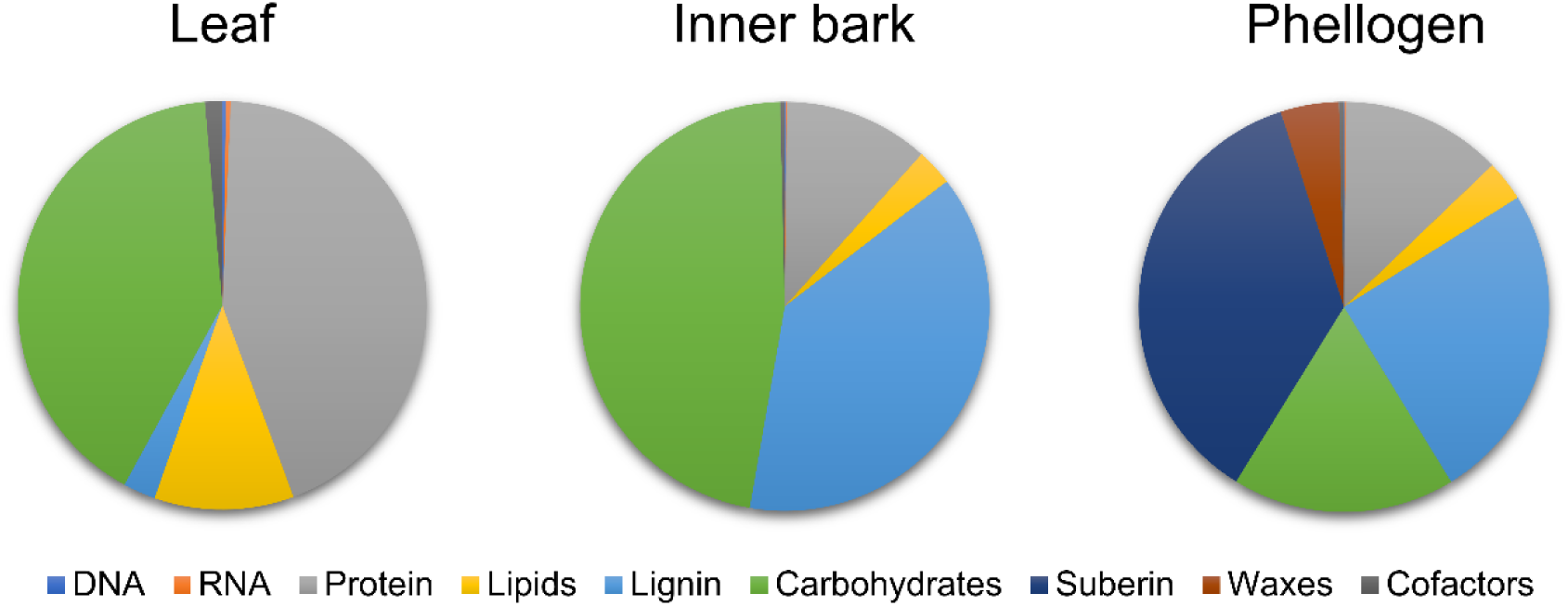
Biomass composition of the leaf, inner bark, and phellogen determined using data retrieved from published plant GSM models and available experimental data. The biomass composition and the respective data sources are detailed in Supplementary File 3.

As explained in the Materials and Methods section, the biomass was formulated by creating “e-Metabolites”, representing the macromolecular composition of each tissue. Each macromolecule (e.g. e-RNA) is associated with a reaction responsible for producing it from its precursors (e.g. ATP, GTP, CTP, UTP).

The leaf macromolecular contents were determined using *A. thaliana* models (Dal’Molin *et al*., 2010; Arnold & Nikoloski, 2014). The cell wall sugar content was included in the e-Carbohydrate composition, while lignin was included in the e-Lignin composition. The monomer contents of DNA, RNA, and protein were determined using the biomass tool available in *merlin*. The fatty acid, lipid, and carbohydrate compositions were determined using experimental data for *Q. suber* or closely related organisms when species-specific data was not available (Koiwai *et al*., 1983; Nouairi *et al*., 2006; Passarinho *et al*., 2006). The lignin, carbohydrate, suberin, and wax contents and composition in the inner bark and phellogen were determined using available experimental data (Pereira, 1988; Lourenço *et al*., 2016).

The Cofactors component includes a set of universal cofactors and vitamins (Xavier *et al*., 2017). These compounds were included in the biomass of each tissue. Nevertheless, the leaf “e-Cofactor” reaction also comprises a set of pigments, such as chlorophylls and carotenoids, determined according to experimental data (Garcia-Plazaola, 1997).

The formulation of tissue-specific biomass composition is critical to obtain tissue-specific models and predict each tissue’s metabolic behavior and the interactions among them. The leaf is mainly composed of carbohydrates and protein. The inner bark also presents high amounts of carbohydrates and a significant content of lignin. The phellogen also exhibits considerable amounts of these macromolecules, but suberin is the one with the highest representation.

For tissue-specific model construction, reactions encoded by genes identified as “not expressed” were removed from each tissue-specific model. The number of genes and reactions decreased in rather different proportions, as *troppo* operates at the reaction level (See Material and Methods section 4.3). Table 3 shows the number of reactions and metabolites in the generic model and each tissue-specific model.

**Table 3.**
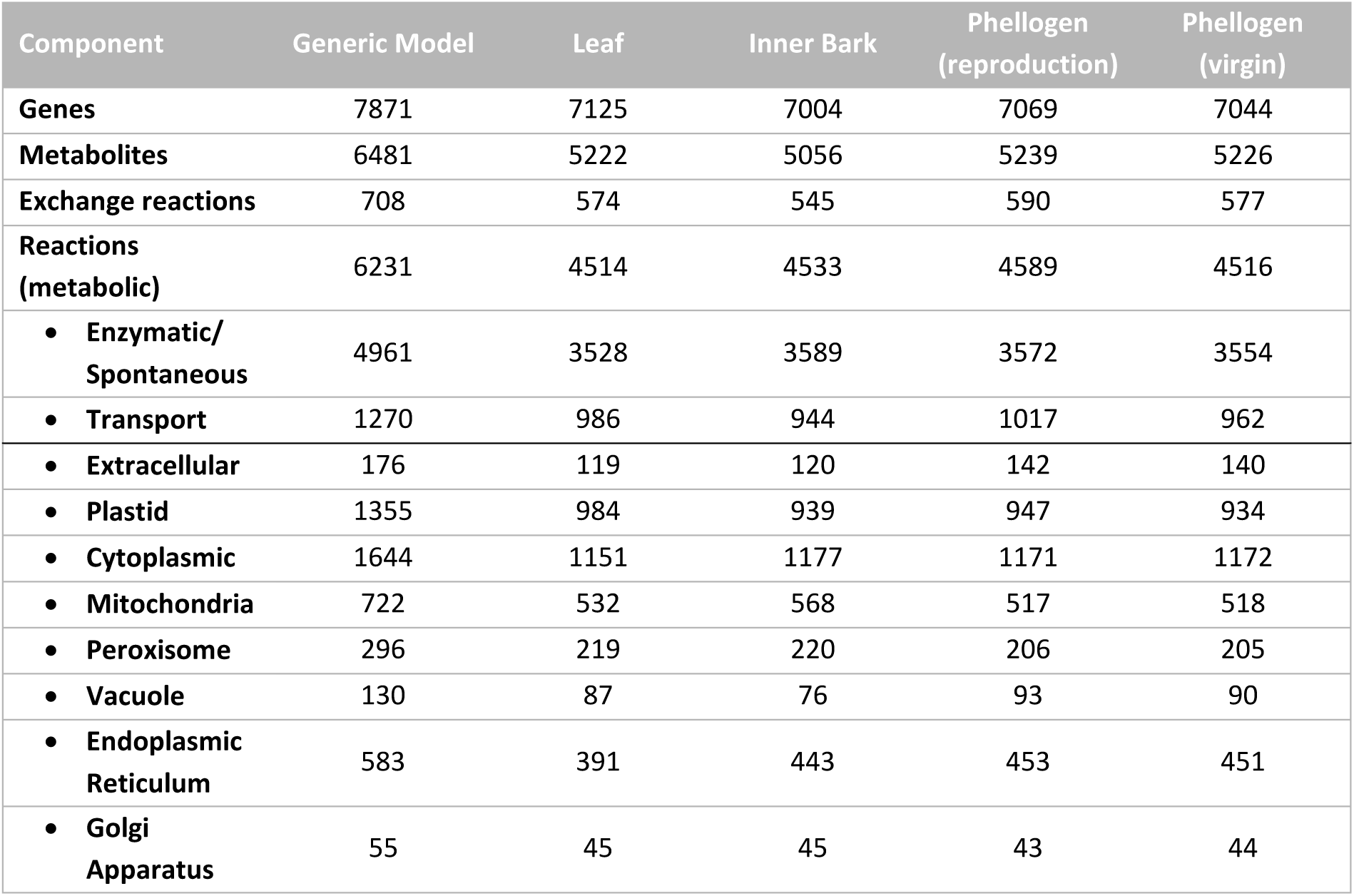
General properties of the generic and tissue-specific GSM models. Genes were predicted based on the cork oak genome and then used to develop the GSM model. The reactions were divided according with the respective metabolic role: metabolic (enzymatic and spontaneous) and transport, and the respective compartment. The number of genes, metabolites, and reactions were determined in the generic model and in each tissue-specific model, generated with *troppo*.

| Component | Generic Model | Leaf | Inner Bark | Phellogen (reproduction) | Phellogen (virgin) |
| --- | --- | --- | --- | --- | --- |
| **Genes** | 7871 | 7125 | 7004 | 7069 | 7044 |
| **Metabolites** | 6481 | 5222 | 5056 | 5239 | 5226 |
| **Exchange reactions** | 708 | 574 | 545 | 590 | 577 |
| **Reactions (metabolic)** | 6231 | 4514 | 4533 | 4589 | 4516 |
| • **Enzymatic/ Spontaneous** | 4961 | 3528 | 3589 | 3572 | 3554 |
| • **Transport** | 1270 | 986 | 944 | 1017 | 962 |
| • **Extracellular** | 176 | 119 | 120 | 142 | 140 |
| • **Plastid** | 1355 | 984 | 939 | 947 | 934 |
| • **Cytoplasmic** | 1644 | 1151 | 1177 | 1171 | 1172 |
| • **Mitochondria** | 722 | 532 | 568 | 517 | 518 |
| • **Peroxisome** | 296 | 219 | 220 | 206 | 205 |
| • **Vacuole** | 130 | 87 | 76 | 93 | 90 |
| • **Endoplasmic Reticulum** | 583 | 391 | 443 | 453 | 451 |
| • **Golgi Apparatus** | 55 | 45 | 45 | 43 | 44 |

The number of metabolic reactions is quite similar among the four tissue-specific models. Nevertheless, the number of reactions in the leaf model’s plastid is slightly higher than in the other models, while in the endoplasmic reticulum, the opposite behavior can be observed. The reproduction phellogen model presents a higher number of transport reactions than the remaining ones. To assess potential differences in the metabolism of the reproduction and virgin phellogen, *in silico* simulations with parsimonious flux balance analysis (pFBA) (Lewis *et al*., 2010) and flux variability analysis (FVA) were performed (Supplementary File 4 – Table S1).

The number of metabolic reactions associated with each pathway is available in Supplementary File 2 Tables S8-S9. The number of reactions associated with the “Arachidonic acid metabolism”, “Brassinosteroid biosynthesis”, and “Drug metabolism – cytochrome P450” pathways are higher in the inner bark and phellogen than in the leaf. The “Cutin, suberin and wax biosynthesis” pathway is mostly represented in the phellogen. As expected, the “Carotenoid biosynthesis” pathway is present in the leaf, while in the remaining tissues, the number of reactions is more limited. No significant differences were identified between the number of reactions in each pathway between the reproduction and virgin phellogen models.

### 2.4. Model Validation

The models developed in this work were tested to guarantee that no biomass or energy is produced without energy input in each condition. The general and tissue-specific metabolic models were validated by analyzing the fluxes of *in silico* simulations in photoautotrophic, heterotrophic, and photorespiratory conditions. The results for model’s validation are available in detail in Supplementary File 5 Tables S1-S6 and Fig. S1-S2. All models are able to grow in heterotrophic conditions, while in photoautotrophic and photorespiratory conditions only the general and leaf models produce biomass.

As expected, the photoautotrophic growth is supported by the assimilation of CO_2_ by RubisCO using light as energy source. The predicted photon uptake flux was 53.68 *mmol* · *gDW*^−1^ · *h*^−1^ for a biomass production fixed at 0.11 h^-1^ (see Materials and Methods). In heterotrophic conditions using sucrose as carbon source, the tricarboxylic cycle (TCA) and the oxidative phosphorylation become more active providing energy to sustain growth. The metabolic response to photorespiration was also assessed in the leaf model. In these conditions, the ‘Glyoxylate and dicarboxylate metabolism’ pathway, which includes the reactions associated with photorespiration, becomes more active to recycle carbon skeletons (Supplementary File 5 Fig. S1).

Quantum yield (amount of CO_2_ fixed per mol of photons) and assimilation quotient (CO_2_ fixed per O_2_ evolved) are common measures of photosynthetic efficiency. Thus, these parameters were determined for the leaf model. In photoautotrophic conditions, the photon yield was calculated as 0.078 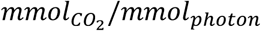, which is within the range reported for *Q. suber* (0.051 – 0.089 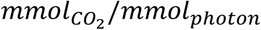) at different light conditions and times of the year (Vaz *et al*., 2010, 2011). Photon yield was also assessed in photorespiration by varying the ratio of carboxylation/oxygenation activity of Rubisco using nitrate and ammonia as nitrogen source (Supplementary File 5 Fig. S2). The results show that the model is able to predict a lower photosynthetic efficiency in drought conditions (carboxylation/oxygenation < 3.0), and when using nitrate as nitrogen source. *In silico* simulations predict an assimilation quotient of 0.76 and 0.93 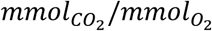 with nitrate and ammonia as nitrogen source, respectively. These values are similar to the ones predicted in other plant leaf GSM models (Poolman *et al*., 2013, 2014).

The inner bark and phellogen metabolic models produce all the metabolites defined in their biomass composition in heterotrophic conditions (Supplementary File 5 Tables S4-S6), using sucrose as carbon source and obtaining energy through the TCA and oxidative phosphorylation. As expected, these models are not able to grow in photoautotrophic conditions as several genes associated with photosynthesis were considered as not expressed by *troppo*, thus, the respective reactions were removed.

### 2.5. Diel Multi-tissue model

A diel multi-tissue metabolic model was generated to analyze the metabolic interactions between leaf, inner bark, and phellogen at the two phases of the diel cycle: light (day) and dark (night). Common pools were created to connect the different tissues for the light and dark phases. The diel multi-tissue GSM model comprises 28,426 reactions and 29,489 metabolites. This model allows the uptake of minerals, such as *HNO*_3_ and *H*_3_*PO*_4_ through the inner bark since the root was not considered. The light/dark uptake ratio of nitrate was constrained to 3:2, as suggested in other diel GSM models (Cheung *et al*., 2014; Shaw & Cheung, 2018). The exchange of oxygen and carbon dioxide is only allowed in the leaves, where photons can be uptake in the light phase. FVA and pFBA were computed, setting as objective function the minimization of photons uptake for photorespiratory conditions. Figure 3 depicts both the relevant transport reactions between tissues, and the metabolites stored between the light and dark phases. The suberin, lignin, and cell wall sugars synthesis pathways are also portrayed. The detailed simulation results are available in Supplementary File 4 – Tables S2-S3.

**Figure 3.**
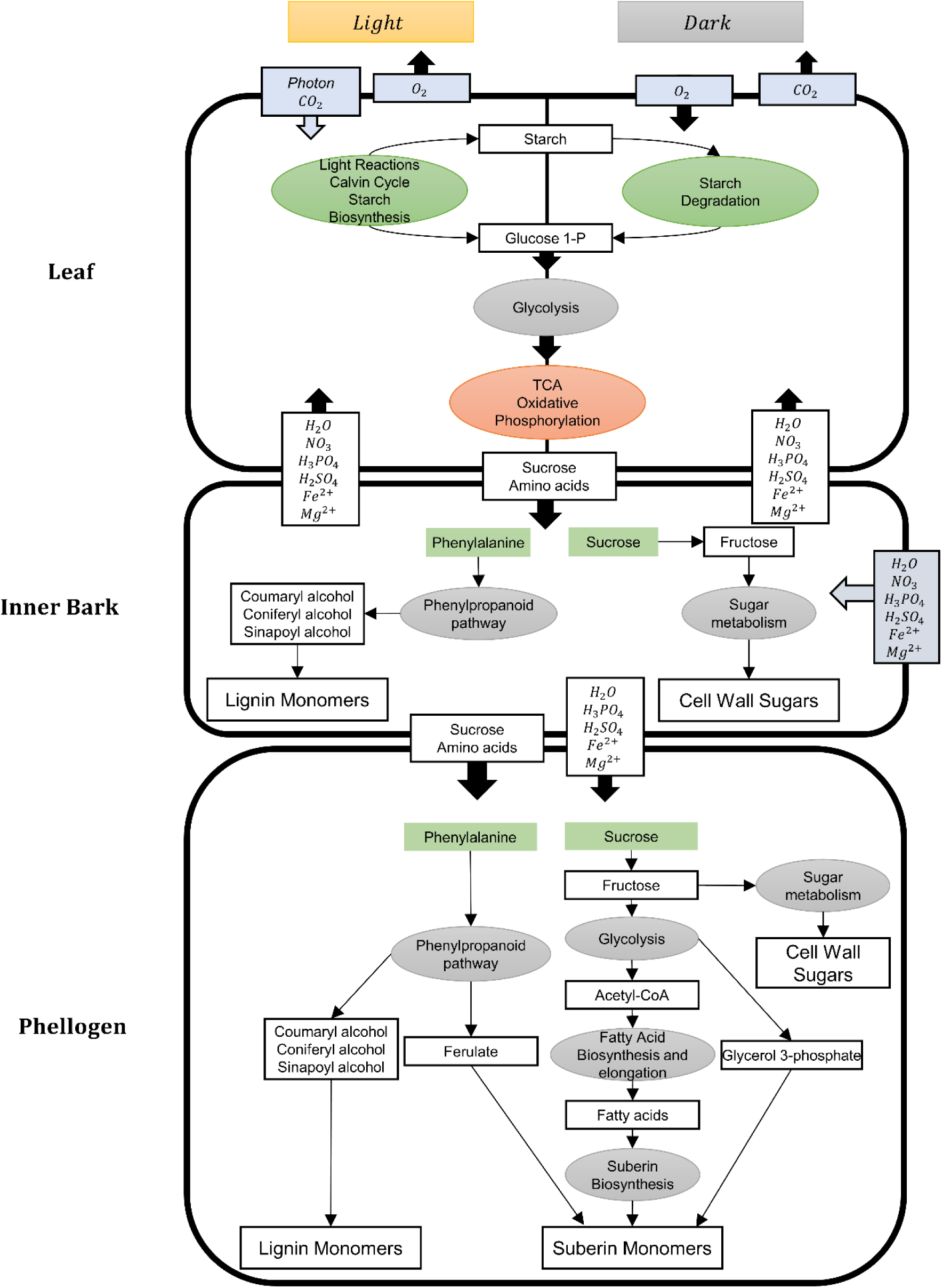
Schematic representation of the metabolic routes towards cork formation. Photons and gases exchanges take place in the leaf, while the uptake of inorganic ions was assumed to happen in the inner bark. Sucrose and amino acids produced in the leaf are transported into the inner bark, and then to the phellogen, where they are used for the suberin and lignin biosynthesis (besides the other biomass components). The differences in the metabolism in the day and night phases are only represented for the leaf since, according with *in silico* simulations, the pathways used in the inner bark and phellogen was similar in the two phases of the diel cycle. The representation is based on pFBA and FVA predictions.

## 3. Discussion

### 3.1. Comparison with other models

*A. thaliana* and *Z. mays* are reference organisms for C3 and C4 plants; thus, several models for these organisms are available. Nevertheless, such models for less studied plants are also becoming available (Grafahrend-Belau *et al*., 2013; Yuan *et al*., 2016b; Pfau *et al*., 2018; Moreira *et al*., 2019; Shaw & Maurice Cheung, 2019). The iEC7871 was compared with *A. thaliana* and *Z. mays* models that provide KEGG identifiers, which facilitates the comparison.

The cork oak model presents 1117 reactions and 945 metabolites not available in the other two models. Interestingly, the annotation of the genes associated with most of these reactions was based on *A. thaliana*’s homologs. The information available in Swiss-Prot for such species is exceptionally high, while this information is scarce for woody plants, which may lead to wrong and missing gene annotations as *Q. suber* is significantly different from *A. thaliana*. Another obstacle is the presence of incomplete EC numbers. For instance, over 300 genes were initially annotated with incomplete EC numbers associated with Cytochromes P450.

The KEGG Pathways associated with these reactions were also analyzed. The pathways with the higher number of unique reactions are associated with the metabolism of xenobiotics, terpenoids, and other secondary metabolites. Also, the “Tryptophan metabolism” and “Tyrosine metabolism” pathways include the secondary metabolism of these amino acids, such as the quinolinic acid production, dopamine, and its derivates.

When classified according to the main areas of the metabolism in KEGG, the pathways representing ‘Lipid metabolism’, ‘aminoacid metabolism and carbohydrate metabolism’, and ‘Xenobiotics biodegradation metabolism’ were the most represented regarding reactions that are only present in iEC7871.

The high number of reactions and metabolites only available in the cork oak model is most likely associated with different reasons: the approach followed to develop the GSM model; the information available at each reconstruction time; the metabolic differences between the three species; the inclusion of pathways associated with secondary plant metabolism.

### 3.2. Secondary Metabolism Pathways

Plants produce a wide range of compounds through their secondary metabolism, whose function includes defense against abiotic and biotic stress or beneficial interactions with other organisms (Isah, 2019). Many secondary metabolites have central roles in the pharmaceutical, cosmetics, perfume, dye, and flavor industries. Despite the recent advances in the investigation of plant secondary metabolism, detailed knowledge of these pathways is restricted to a few species (Isah, 2019), such as *A. thaliana, O. sativa*, and *Z. mays*. These pathways are not always complete or available in biological databases, implying an additional effort to find information regarding genes, reactions, and metabolites associated with secondary plant metabolism. Although there is still much to learn about these compounds and respective biosynthetic pathways, genome-scale modelling can provide insights into a given organism’s potential to produce secondary metabolites. The cork oak model reconstructed in this work includes reactions associated with the biosynthesis and metabolism of several secondary metabolites, steroids, and drugs.

The “Metabolism of xenobiotics by cytochrome P450” includes 38 reactions not found in the *A. thaliana* and *Z. mays* models considered in the Results section. This pathway represents a set of reactions, mostly associated with the cytochrome P450, responsible for the response to the presence of toxic xenobiotics. The model includes the complete routes for the degradation of benzopyrene, aflatoxin B1, and nicotine-derived xenobiotics. The cytochrome P450 family has been described being highly upregulated in developing phellem tissues (Lopes *et al*., 2020).

The model presents all the necessary reactions to produce jasmonic acid and its derivates, usually called jasmonates, through the “alpha-Linoleic acid metabolism” pathway. These plant hormones are associated with the regulation of growth and developmental processes, stomatal opening, inhibition of Rubisco biosynthesis, and nitrogen and phosphorus uptake (Wasternack & Hause, 2013; Ruan *et al*., 2019). The “Steroid biosynthesis” pathway is essentially complete, enabling the production of steroids like ergosterol, cholesterol, and stigmasterol. The enzymes associated with the synthesis of calcitetrol and secalciferol (animal hormones) are the only ones whose encoding genes were not found in the genome of *Q. suber*. Although these hormones are usually synthesized by animals, they were already identified in a few plants (Aburjai *et al*., 1998; Boland *et al*., 2003; Jäpelt & Jakobsen, 2013). The biosynthetic pathway for the synthesis of gibberellins is also available in the model. Although the regulatory effect of hormones and steroids cannot be quantitively evaluated in stoichiometric models, the presence of their biosynthetic pathways, connected to the core network, can be used as a source of information regarding the potential to produce a certain hormone or steroid.

Although this model contains a considerable amount of information regarding secondary metabolism, further curation of these pathways would improve the connectivity of the model and reduce the existing gaps.

### 3.3. Suberin, Lignin, and Waxes Biosynthesis

As mentioned before, suberin is the major component of phellogen, while its abundance is residual in the leaf and inner bark (Figure 2). The phellogen also contains significant amounts of lignin and waxes. The biosynthesis pathway of the monomers of these cork components in the model is represented in Figure 4.

**Figure 4.**
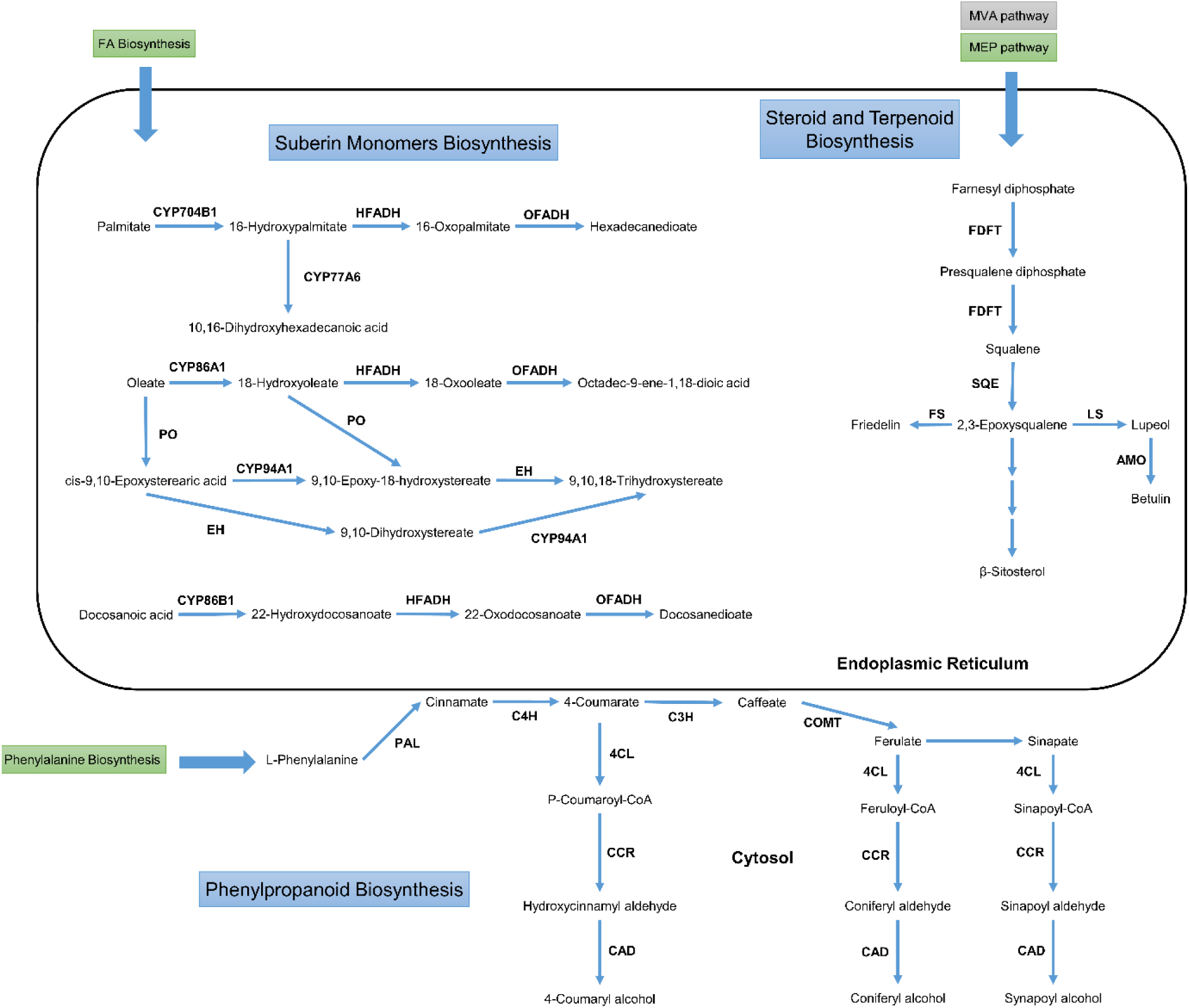
Biosynthetic pathway of suberin, waxes, and lignin monomers. The C:16 and C:18 fatty acids produced in the plastid are transported to the endoplasmic reticulum, where they can be elongated and unsaturated, and follow the suberin monomer biosynthesis pathway. Cytochromes P450 (CYPs), peroxygenases, epoxy hydroxylases, ω-oxo-fatty acid dehydrogenases, ω-hydroxy-fatty acid dehydrogenases catalyze successive reactions to produce the aliphatic monomers. Farnesyl diphosphate, produced from isopentenyl diphosphate provided by the cytosolic mevalonate (MVA) pathway or by the plastidic methylerythritol phosphate (EMP) pathway is used as the initial precursor of steroids in the ER. 2,3-Epoxysqualene is converted into sterols and terpenoids, monomers of the wax component of the phellogen. The phenylpropanoid pathway uses phenylalanine produced in the leaf’s chloroplasts, to produce cinnamate. The pathway follows in the cytosolic surface of the endoplasmic reticulum, and then in the cytosol, producing the lignin monomers. 4CL – 4-Coumarate CoA ligase, AMO: β-amyrin monooxygenase, C3H: p-coumarate 3-hydroxylase, C4H: cinnamate 4-hydroxylase, CAD: cinnamyl alcohol dehydrogenase, CCR: cinnamoyl-CoA reductase, COMT: Caffeoyl-CoA O-methyltransferase, EH: epoxide hydroxylase, FDFT: farnesyl diphosphate farnesyltransferase, FS: friedelin synthase, HFADH: ω-hydroxy-fatty acid dehydrogenase, LS - lupeol synthase, OFADH: ω-oxo-fatty acid dehydrogenase, PAL: phenylalanine ammonia lyase, PO: peroxygenase, SQE: squalene monooxygenase. The IDs of the reactions catalyzed by these enzymes can be found in Table S10 of Supplementary File 2.

The synthesis of the aliphatic suberin monomers is associated with the KEGG’s “Cutin, suberin, and wax biosynthesis” pathway. Fatty acids are exported from the plastid and imported into the endoplasmic reticulum, where these can be elongated to originate long-chain and very long-chain fatty acids. Cytochromes P450, especially of the CYP86A family, are responsible for catalyzing several reactions that synthesize suberin monomers. CYP86A1 and CYP704B1 are responsible for the ω-hydroxylation of long-chain fatty acids, while CYP86B1 acts on very-long chain fatty acids.

Epoxide hydrolases (EC: 3.3.2.10), ω-oxo-fatty acid dehydrogenases (EC: 1.2.1.-), ω-hydroxy-fatty acid dehydrogenases (EC: 1.1.1.-), and peroxygenases (EC: 1.11.2.3) can act in the ω-hydroxy acid in different combinations, to originate the diverse α,ω-dicarboxylic, poly α,ω-dicarboxylic, and polyhydroxy-fatty acids. The genome annotation and the model manual curation allowed identifying 38 genes encoding epoxide hydrolase, five encoding ω-hydroxy-fatty acid dehydrogenase, and six encoding peroxygenases. Nevertheless, no genes encoding ω-oxo-fatty acid dehydrogenase were available in any database; thus, the reactions catalyzed by this enzyme were added without gene association.

The polymerization process is still not completely understood but it likely involves esterification reactions through glycerol-3-phosphate acyltransferases (GPAT), producing acylglycerol esters. This family of enzymes was identified as a key step in suberin/cork polymerization and the expression level of GPAT5 gene is higher in June, which corresponds to a period of higher phellogen activity (Marum *et al*., 2011). Acylglycerol esters are then secreted through the Golgi secretory pathway and ABC transporters (which are overexpressed in phellem (Lopes *et al*., 2020) and incorporated in the suberin glycerol polyester (Vishwanath *et al*., 2015). The aliphatic suberin polyester transport and assembly were simplified in the model and represented through a reaction converting the determined precursors (Supplementary File 3 – Table S6) into the macromolecular representation of suberin: “e-Suberin”.

Waxes are also an important cork component and are composed essentially of sterols and terpenoids (Castola *et al*., 2005). These are produced through the “Steroid biosynthesis” pathway in the endoplasmic reticulum. The initial precursor, farnesyl diphosphate, is synthesized in the cytosolic mevalonate pathway or by the plastidic methylerythritol phosphate pathway. Farnesyl diphosphate farnesyltransferase and squalene monooxygenase convert farnesyl diphosphate into 2,3 – epoxysqualene. This triterpenoid can be converted into cyclic tripterpenoids, such as friedelin and lupeol, or can be used to produce phytosterols like β-sitosterol, through a longer pathway.

The cork contains a considerable amount of lignin, although this polymer is present in other tissues. The synthesis of the respective monomers (guaiacyl lignin, hydroxyphenyl lignin, and syringyl lignin) is described in the KEGG’s “Phenylpropanoid biosynthesis” pathway. Phenylalanine is the main precursor of this pathway that occurs mostly in the cytoplasm. It is converted to cinnamic acid by phenylalanine ammonia-ligase (EC: 4.3.1.24). Cinnamate monooxygenase, caffeate methyltransferase, and ferulate-5-hydroxylase are responsible for the successive conversion of cinnamic acid to coumarate, ferulate, hydroxyferulate, and sinapic acid, in the cytosolic surface of the endoplasmic reticulum. These metabolites can be converted into lignin monomers in a three-step process catalyzed by coumarate-CoA ligase (EC: 6.2.1.12), cinnamoyl-CoA reductase (EC: 1.2.1.44), and cinnamyl-alcohol dehydrogenase (EC: 1.1.1.195). Through the polymer assembly in the cell wall, peroxidases (EC: 1.11.1.7) convert the guaiacyl, hydroxyphenyl, and syringyl alcohols into guaiacyl, hydroxyphenyl, and syringyl lignins, respectively. The linkages between aliphatic and aromatic suberin were not included in the model since these macromolecules are represented in the biomass separately.

### 3.4. Tissue-specific models

A generic metabolic model comprises all the reactions catalyzed by all the enzymes encoded in an organism’s genome. However, this approach does not account for the regulatory network present in each organ, tissue, or cell in different environmental conditions. The integration of transcriptomics data in a GSM model allows obtaining metabolic models closer to the *in vivo* phenotype of the respective tissue or condition.

Transcriptomics data for the leaf, inner bark, and phellogen were integrated into the generic model, originating three tissue-specific models. The biomass of each tissue was determined using data available in the literature and other metabolic models. The leaf macromolecular composition was mostly based on previously published GSM models. Quantitative information regarding the leaf composition of *Q. suber* would be useful to increase the model’s reliability and improve predictions. The composition of the inner bark and the phellogen was based on experimental data available for *Q. suber*. While the inner bark is mostly composed of lignin and carbohydrates, the phellogen also comprises suberin and waxes. Suberin is composed of glycerol, ferulic acid, and diverse alkanols, fatty acids, hydroxy acids, ω-hydroxy acids and α,ω-dicarboxylic acids. Waxes are composed of terpenes, such as friedelin, and sterols, like β-sitosterol. Although the cork bark contains tannins (mostly ellagitannins) (Cadahía *et al*., 1998), these compounds were not included in the biomass since their biosynthetic pathways are poorly understood and not available in the used biological databases.

The number of reactions and metabolites on each subcellular compartment was compared across the three tissues. The number of transport reactions in the Reproduction Phellogen is visibly higher than in the other tissues, including the Virgin Phellogen. These reactions are essentially associated with the transport of steroids and hormones. Based on BLAST searches against TCDB, the genes associated with these reactions are auxin (phytohormone required for cell division (Perrot-Rechenmann & Napier, 2005)) efflux pumps. An analysis of the pathways available in the Reproduction and Virgin Phellogen models, followed by *in silico* simulations did not allow to identify significant differences between them. The cork oak phellogen’s transcriptional profile allows inferring that the flavonoid route is favored in bad quality cork, while the lignin and suberin production pathways are preferred in good quality cork production (Teixeira *et al*., 2018). A recent study reported a higher expression of genes associated with fatty acid biosynthesis and elongation in reproduction cork (Lopes *et al*., 2020). Although it was reported that the macromolecular composition of reproduction and virgin cork could be similar, the precursors of suberin and extractives are significantly different (Pereira, 1988). Hence, a biomass composition specific to the virgin phellogen should be defined, including the tannin content, to allow the observation of metabolic differences between the two models.

### 3.5. Diel multi-tissue model

Tissues and organs of multicellular organisms do not perform their metabolic functions individually. Instead, they interact with each other by exchanging sugars, amino acids, hormones, and others. In plants, the simplest example is the sucrose formation in the leaves and its transport across all other organism tissues. Multi-tissue GSM models allow analyzing the dependencies between tissues in terms of biomass precursors, carbon skeletons, nitrogen, sulfur and phosphorus sources, and the energy required for their translocation and proton balance (Bordbar *et al*., 2011; Zakhartsev *et al*., 2016). The introduction of light and dark phases allows predicting features that are not possible in continuous light models, such as the accumulation and utilization of carboxylic acids (Cheung *et al*., 2014).

The leaf, inner bark, and phellogen models were merged into a multi-tissue GSM model. Light and dark phases were considered to present the metabolic differences that occur in these two phases. Other tissues or organs, such as the root and xylem, could also be considered and included in the multi-tissue model. Nevertheless, this work’s scope is related to the metabolism of cork precursors and including more tissues would increase the complexity of the model unnecessarily.

The most significant differences between the light and dark phases of metabolism were identified in the leaf (Supplementary File 4 – Tables S8-S9), due to the photosynthesis and photorespiration pathways.

In the light dependent reactions, carbon dioxide is fixed through the Calvin Cycle, forming carbohydrates, and consequently, the carbon skeletons for the synthesis of the remaining biomass components. A fraction of the carbohydrates produced here (mainly starch) is stored and used as energy source when needed. The storage of starch is more efficient in the model than sucrose, glucose, or fructose. Whereas starch is mobilized in plastids, soluble sugars are accumulated in the vacuole, implying an energetic cost to its transport. The model produced malate in the dark (light independent reactions), which is used at day (light dependent) by the malic enzyme to produce pyruvate and obtain reducing power (NADPH). Citrate was also produced at night (light independent reaction) through the tricarboxylic cycle (TCA) and used to feed the diurnal TCA and provide carbon skeletons for the amino acid metabolism. This behavior is in agreement with the previously published diel GSM model for *A. thaliana* (Cheung *et al*., 2014).

Nitrate provided by the inner bark is transported into the leaf and is then converted into ammonia. In the plastids, ammonia is used to incorporate nitrogen through the glutamine synthetase-glutamate synthase pathway.

Sucrose and amino acids are exported from the leaf and then imported by proton symport by the inner bark. The proton balance of the common pool is maintained by plasma membrane ATPases. In the inner bark, sucrose is used as an energy and carbon source to produce the biomass components. As mentioned before, the inner bark biomass is mainly composed of carbohydrates, lignin, and protein. Hence, pathways associated with the synthesis of cell wall precursors (“Amino sugar and amino nucleotide metabolism”, “Starch and sucrose metabolism”, and “Phenylpropanoid biosynthesis”) are the most relevant in this tissue.

The remaining sucrose and amino acids produced in the leaves are transported to the phellogen. The reversible sucrose synthase activity allows sucrose conversion into UDP-glucose, which is used to produce other cell wall components, and fructose that follows the glycolytic pathway to produce pyruvate. A fraction of it is transported into the plastid and converted by pyruvate dehydrogenase into acetyl-CoA, the fatty acids precursor. As mentioned above, fatty acids are elongated and suffer a series of hydroxylation, epoxidation, and peroxidation reactions in the endoplasmic reticulum. Ferulate, another precursor of suberin, is produced through the “Phenylpropanoid biosynthesis” pathway, with phenylalanine serving as a precursor. The glycerol 3-phosphate used for the esterification reactions is produced from glycerone phosphate produced through glycolysis.

The diel multi-tissue GSM model developed in this work is a useful framework capable of providing insights into the metabolism of *Q. suber* at a global level. It can be used to study the biosynthetic pathways of suberin, lignin, waxes, and many other compounds.

## Conclusions

In this work, we present the first genome-scale metabolic model of a woody plant. The iEC7871 was based on knowledge retrieved from genomic information, biological databases, and literature. It can simulate the Cork Oak tree’s metabolic behavior in phototrophic and heterotrophic conditions, as well as photorespiration. This model comprises the pathways of the central metabolism and several pathways associated with the secondary metabolism reproducing the formation of the major components of cork.

The integration of transcriptomics data allowed obtaining tissue-specific models for the leaf, inner bark, and phellogen. These models were merged to obtain a multi-tissue GSM model that comprises the diel cycle’s dark and light phases.

This GSM model comprehends the four main secondary metabolic pathways participating in cork production: acyl-lipids, phenylpropanoids, isoprenoids, and flavonoids. The lipid biosynthesis pathway is required for the biosynthesis of the linear long-chain compounds forming the aliphatic suberin domain, which share upstream reactions with waxes biosynthesis. The phenylpropanoid metabolism is needed for the biosynthesis of the cork aromatic components, which share reactions with wood lignin.

The metabolic models developed in this work can be used as a tool to analyze and predict the metabolic behavior of the tree and evaluate its metabolic potential. Metabolic modelling methods can be applied, including dynamic approaches, to study the changes in this tree’s metabolism over time and environment.

## 4. Materials and Methods

### 4.1. Software

*merlin* v4 was used to support the reconstruction process, while COBRApy v0.20.0 (Ebrahim *et al*., 2013) was used to perform all simulations and analyses of the GSM model, as well as generate the diel multi-tissue model. The simulations were performed using the CPLEX v128.0.0 solver.

The Troppo (Ferreira *et al*., 2020) python package was used to integrate the transcriptomics data in the metabolic model, originating tissue-specific models.

FastQC (https://www.bioinformatics.babraham.ac.uk/projects/fastqc/), Sickle (Joshi & Fass, 2011), Bowtie2 (Langmead & Salzberg, 2012), FeatureCounts (Liao *et al*., 2014), and edgeR (Robinson *et al*., 2009) were used to process the transcriptomics data.

### 4.2. Metabolic Reconstruction

The “Automatic workflow” tool, available in *merlin*, was used to perform the genome annotation by assigning EC numbers to enzyme encoding genes based on BLAST searches against Swiss-Prot and TrEMBL.

A draft metabolic network was assembled by loading KEGG’s metabolic information and integrating the genome annotation. KEGG reactions associated with enzymes identified in the genome annotation stage were included in the model, as well as spontaneous reactions.

Transport reactions were automatically generated using the TranSyT (Lagoa *et al*., 2021) tool. Nevertheless, additional transport reactions were added if reported in the literature, or if necessary for the model functionality. The subcellular location of proteins was predicted using the WolfPsort tool. Additionally, LocTree3 (Goldberg *et al*., 2014) and ChloroP 1.1 (Emanuelsson *et al*., 1999) were used to verify protein locations during the manual curation stage.

The leaf, inner bark, and phellogen biomass compositions were based on previously published plant GSMMs and available literature. The biomass precursors were organized in macromolecules or cell structures, labelled as “e-Metabolites”. The leaf macromolecular contents were determined using *A. thaliana* models (Dal’Molin *et al*., 2010; Arnold & Nikoloski, 2014). The cell wall sugar content was included in the e-Carbohydrate composition, while lignin was included in the e-Lignin composition. The monomer contents of DNA, RNA, and protein were determined using the biomass tool, available in *merlin*. The fatty acid, lipid, and carbohydrate compositions were determined using experimental data for *Q. suber* or closely related organisms when species-specific data were not available (Koiwai *et al*., 1983; Nouairi *et al*., 2006; Passarinho *et al*., 2006). The lignin, carbohydrate, suberin, and wax contents and composition in the inner bark and phellogen were determined using available experimental data (Pereira, 1988; Lourenço *et al*., 2016). Suberin was only accounted for in the inner bark and phellogen. The e-Cofactor component includes a set of universal cofactors and vitamins (Xavier *et al*., 2017), which were included in the biomass of each tissue. Nevertheless, the leaf “e-Cofactor” reaction also comprises a set of pigments, such as chlorophylls and carotenoids, determined according to experimental data (Garcia-Plazaola, 1997). The energy requirements were inferred as reported by (Yuan *et al*., 2016a).

Through the whole process, literature, and biological databases, namely KEGG, MetaCyc (Caspi *et al*., 2020), and BRENDA (Chang *et al*., 2021), were consulted to retrieve information regarding metabolic, genomic, and enzymatic information, as described previously (Thiele & Palsson, 2010; Dias *et al*., 2019). Briefly, KEGG Pathways allowed identifying reactions that were not included through the automatic network assembly; MetaCyc was used to identify reactions not available in KEGG; MetaCyc and BRENDA provided information regarding the reaction’s reversibility and mass-balance.

The validation of the GSM model started by assuring the biomass formation, which was performed using the BioISO plug-in available in *merlin*.

The following validation approaches were applied to the model:

1. Growth without photons, carbon, nitrogen, phosphorus, and sulfur sources.
2. Futile cycles and stoichiometrically balanced cycles.
3. Growth rate assessment.
4. Growth with different elemental sources.
5. Capacity to present flux through photosynthesis, respiration, and photorespiration.

The model validation was performed using FBA (Orth *et al*., 2010), pFBA, and FVA, available in COBRApy. The general GSM model of *Q. suber* is available in SBML format in Supplementary File 6. All the metabolic models developed in this work are available at https://cutt.ly/quercussubermodels.

### 4.3. Tissue-Specific and multi-tissue Models

Transcriptomics data from different tissues were used to obtain tissue-specific models. The quality of the transcriptomics data was analyzed using the FastQC software. After trimming the data with Sickle, Bowtie2 was used to align the trimmed FASTQ files against the reference genome. FeatureCounts was used to count the mapped reads to each gene. After filtering and normalizing the data with the edgeR library, datasets with the normalized counts of each gene were obtained. These files were used as input for *troppo*, together with the generic GSMM, to generate tissue-specific models. *Troppo* was run using the CPLEX solver and the fastcore reconstruction algorithm. A *troppo*’s integration strategy was developed to guarantee that the tissue-specific models are capable of producing biomass. The median of each dataset was used as threshold, and the remaining parameters were kept as default. *Troppo* identifies the reactions that should be removed from the generic model, based on the expression of the genes included in the gene-protein-reaction association of each reaction. Nevertheless, the reactions that were maintained in the model within this approach kept all of their respective genes. The carotenoid biosynthesis has two branches, one associated with trans-Phytofluene and the other with 15,9-dicis-Phytofluene. The enzymes present in the two branches are similar (phytoene desaturase and zeta-carotene desaturase). However, one of them is dependent on photosynthesis since it unbalances the plastoquinone/plastoquinol ratio. The other one is stoichiometrically balanced since it involves successive oxidation and reduction of these quinones. Hence, the presence of the second branch was guaranteed in the leaf model since it is essential for the carotenoid production through the dark phase.

The tissue-specific models obtained using *troppo* were adapted to account for the light and dark conditions, by duplicating all reactions and metabolites. Reactions converting sugars (starch, sucrose, glucose, fructose, malate, citrate, quercitol, and quinate), 18 amino acids, and nitrate between light and dark were added to the model, based on a previously published approach (Cheung *et al*., 2014). To build the multi-tissue model, the diel tissue-specific models were merged, and common pools between leaf and inner bark (common pool 1), and inner bark and phellogen (common pool 2) were created. Since the root was not accounted for, the uptake of water, nitrogen, sulfur, phosphorus, and magnesium sources was included in the inner bark.

### 4.4. Simulations

Two different approaches were used to perform simulations with pFBA. The first was to fix the biomass formation to 0.11 *h*^−1^, and set the minimization of the photon uptake as the objective function. In the second approach, the photon uptake was fixed to 100 *mmol*_*photon*_. *gDW*^−1^. *h*^−1^, maximizing the biomass production. Photorespiration was simulated by constraining the carboxylation/oxygenation flux ratio (Vc/Vo) of Rubisco (reactions R00024 and R03140) to 3:1 (Weber, 2007). Simulations with the inner bark, and virgin and reproduction phellogen were performed by maximizing the biomass production while limiting the sucrose and amino acid uptake to 1 *mmol*_*photon*_. *gDW*^−1^. *h*^−1^.

The simulations with the diel multi-tissue GSM model were performed using the restrictions applied in photorespiration, with the additional constraint of the nitrate light/dark uptake ratio, which was settled to 3:2 (Cheung *et al*., 2014; Shaw & Cheung, 2018).

### 4.5. Accession Numbers

The genome sequence of *Quercus suber* was retrieved from the NCBI database with the assembly accession number GCF_002906115.1. The transcriptomics data was retrieved from the EBI database(Li *et al*., 2015) (Accession PRJNA392919 for Leaf and Inner Bark; Accession PRJEB33874 for Phellogen).

## 5. Supporting Information

5.1. Supplementary file 1: Results of the genome annotation, and respective analysis.

5.2. Supplementary file 2: Reactions and metabolites with KEGG IDs, included in the cork oak, A. thaliana, and Z. mays models (S1-S5). Analysis of the pathways whose reactions were only identified in the cork oak model according with the genome annotation (S6), and by tissue (S7). Distribution of reactions by pathway in each tissue (S8).

5.3. Supplementary file 3: Biomass Composition. Data collected from publications, and respective treatment to include it in the GSM models.

5.4. Supplementary file 4: Flux values, maximum, and minimum according to with pFBA and FVA simulations for the reproduction and virgin phellogen models, and the transport reactions across tissues and storage reactions in the diel multi-tissue model.

5.5. Supplementary file 5: Validation of the metabolic models.

5.6. Supplementary file 6: iEC7871 model in SBML format.

## 6. Acknowledgments and Funding

The authors would like to acknowledge project 22231/01/SAICT/2016: “Biodata.pt – Infraestrutura Portuguesa de Dados Biológicos”, supported by Lisboa Portugal Regional Operational Programme (Lisboa2020), under the PORTUGAL 2020 Partnership Agreement, through the European Regional Development Fund (ERDF). The authors would also like to acknowledge the Portuguese Foundation for Science and Technology (FCT) for the Strategic Funding FCT 2020-2023 (PEst UIDB/04469/2020). E. Cunha acknowledges FCT for the scholarship (DFA/BD/8076/2020). Oscar Dias acknowledges FCT for the Assistant Research contract obtained under CEEC Individual 2018. Inês Chaves was funded by DL 57/2016/CP1351/CT0003.

## 7. Author Contributions

E.C. and M.S. developed and tested the metabolic models; I.C. analyzed the data regarding biochemical and molecular biology aspects of the cork oak; H.D. and D.L. processed the transcriptomics data; D.R.L. developed methods for data processing and analysis, and model reconstruction; E.C. wrote the original draft; I.C., M.R., I.R., and O.D. reviewed and edited the article; M.R., I.R., and O.D. conceived and supervised the project.

## 8. Data Availability

All metabolic models developed in this work are available at https://bit.ly/38Iez8h.

