## Supplementary File 5 for "The first multi-tissue diel genome-scale metabolic model of a woody plant highlights suberin biosynthesis pathways in *Quercus suber*"

Table S1. Summary of a pFBA applied to the leaf model in photoautotrophic conditions. The Biomass was fixed to 0.11 h^-1^ and the objective function was defined as the minimization of photon uptake.

| Reaction | Metabolite | Flux |
| --- | --- | --- |
| EX_C00205__dra | hn | -53.674812 |
| EX_C00011__dra | CO2 | -4.236668 |
| EX_C00001__dra | H2O | -2.817708 |
| EX_C00244__dra | Nitrate | -0.520394 |
| EX_C00059__dra | Sulfate | -0.016141 |
| EX_C00009__dra | Orthophosphate | -0.007504 |
| EX_C00080__dra | H+ | -5e-06 |
| EX_C00305__dra | Magnesium cation | -2e-06 |
| EX_C14818__dra | Fe2+ | -1e-06 |
| EX_C00237__dra | CO | 2e-06 |
| EX_Biomass__cyto | e-Biomass | 0.11 |
| EX_C00007__dra | Oxygen | 5.582437 |

Table S2. Summary of a pFBA applied to the leaf model in heterotrophic conditions. The Biomass was fixed to 0.11 h^-1^ and the objective function was defined as the minimization of sucrose uptake.

| Reaction | Metabolite | Flux |
| --- | --- | --- |
| EX_C00007__dra | Oxygen | -1.810577 |
| EX_C00089__dra | Sucrose | -0.616084 |
| EX_C00244__dra | Nitrate | -0.520394 |
| EX_C00059__dra | Sulfate | -0.016141 |
| EX_C00009__dra | Orthophosphate | -0.007504 |
| EX_C00305__dra | Magnesium cation | -2e-06 |
| EX_C14818__dra | Fe2+ | -1e-06 |
| EX_C00237__dra | CO | 2e-06 |
| EX_C00288__dra | HCO3- | 5e-06 |
| EX_Biomass__cyto | e-Biomass | 0.11 |
| EX_C00011__dra | CO2 | 3.156341 |
| EX_C00001__dra | H2O | 3.959216 |

Table S3. Summary of a pFBA applied to the leaf model in photorespiratory conditions. The biomass was fixed to 0.11 h^-1^ and the objective function was defined as the minimization of photon uptake. The carboxylation/oxygenation ratio of Rubisco was fixed to 3:1.

| Reaction | Metabolite | Flux |
| --- | --- | --- |
| EX_C00205__dra | hn | -74.164666 |
| EX_C00011__dra | CO2 | -4.236668 |
| EX_C00001__dra | H2O | -2.817708 |
| EX_C00244__dra | Nitrate | -0.520394 |
| EX_C00059__dra | Sulfate | -0.016141 |
| EX_C00009__dra | Orthophosphate | -0.007504 |
| EX_C00080__dra | H+ | -5e-06 |
| EX_C00305__dra | Magnesium cation | -2e-06 |
| EX_C14818__dra | Fe2+ | -1e-06 |
| EX_C00237__dra | CO | 2e-06 |
| EX_Biomass__cyto | e-Biomass | 0.11 |
| EX_C00007__dra | Oxygen | 5.582437 |

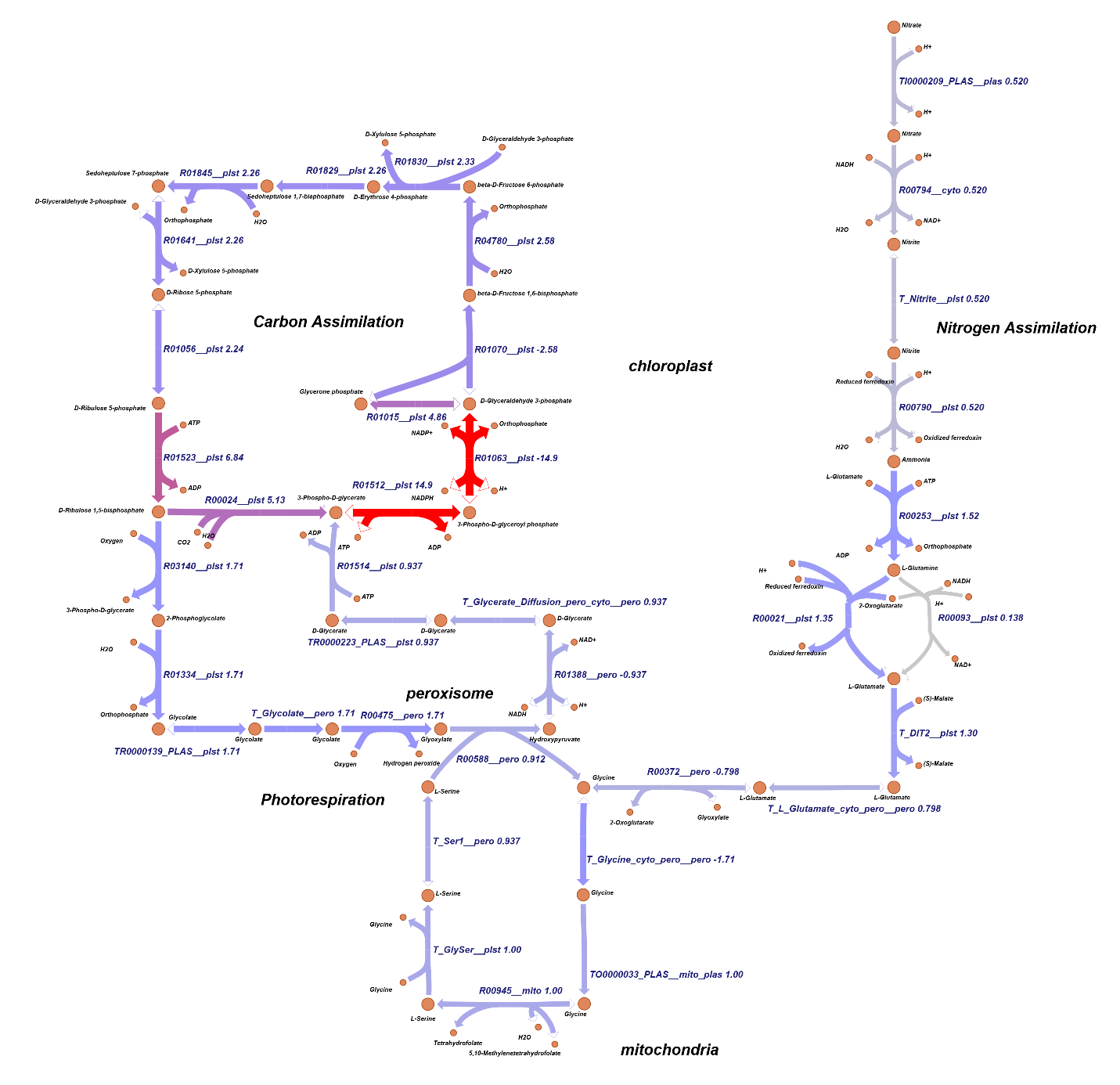

Figure S1. Metabolic map of carbon and nitrogen assimilation, and photorespiratory pathway during photorespiration based on the simulation mentioned in Table S3.

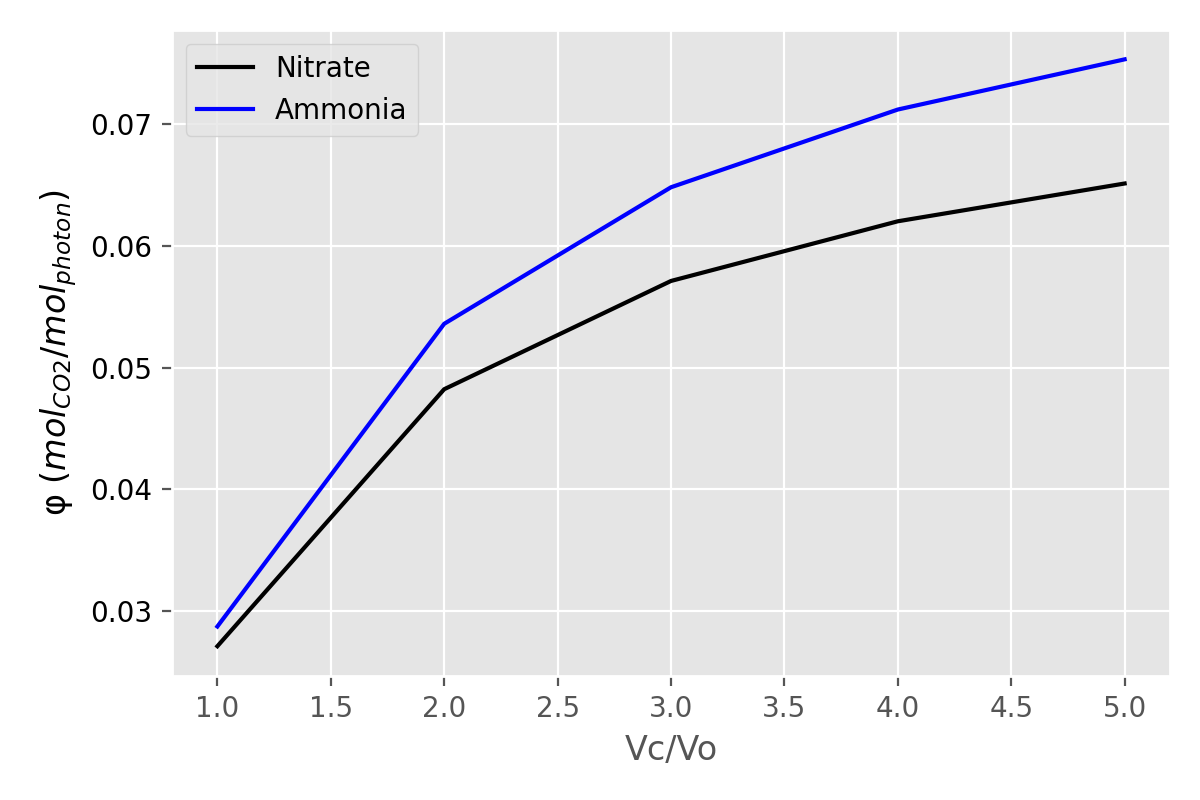

Figure S2. Variation of the quantum yield (mol CO_2_ fixed per mol of photon) with the carboxylation/oxygenation ratio (Vc/Vo) of Rubisco. The biomass was fixed to 0.11 h^-1^ and the objective function was defined as the minimization of photon uptake.

Table S4. Summary of a pFBA applied to the inner bark in heterotrophic conditions. The maximum uptake of amino acids and sucrose was fixed to 1 $mmol\cdot g_{DW}\cdot h^{-1}$, while maximizing the biomass production

| Reaction | Metabolite | Flux |
| --- | --- | --- |
| EX_C00007__dra | Oxygen | -20.508113 |
| EX_C00049__dra | L-Aspartate | -1.0 |
| EX_C00062__dra | L-Arginine | -1.0 |
| EX_C00082__dra | L-Tyrosine | -1.0 |
| EX_C00079__dra | L-Phenylalanine | -1.0 |
| EX_C00078__dra | L-Tryptophan | -1.0 |
| EX_C00152__dra | L-Asparagine | -1.0 |
| EX_C00065__dra | L-Serine | -1.0 |
| EX_C00064__dra | L-Glutamine | -1.0 |
| EX_C00089__dra | Sucrose | -1.0 |
| EX_C00041__dra | L-Alanine | -1.0 |
| EX_C00037__dra | Glycine | -1.0 |
| EX_C00025__dra | L-Glutamate | -1.0 |
| EX_C00188__dra | L-Threonine | -1.0 |
| EX_C00123__dra | L-Leucine | -0.07058 |
| EX_C00183__dra | L-Valine | -0.04705 |
| EX_C00047__dra | L-Lysine | -0.041571 |
| EX_C00407__dra | L-Isoleucine | -0.037902 |
| EX_C00135__dra | L-Histidine | -0.019391 |
| EX_C00073__dra | L-Methionine | -0.018013 |
| EX_C00009__dra | Orthophosphate | -0.015 |
| EX_C00059__dra | Sulfate | -0.014254 |
| EX_C14818__dra | Fe2+ | -2e-06 |
| EX_C00237__dra | CO | 3e-06 |
| EX_C00288__dra | HCO3- | 1.4e-05 |
| EX_Biomass__cyto | e-Biomass | 0.837534 |
| EX_C00086__dra | Urea | 0.954851 |
| EX_C00628__dra | 2,5-Dihydroxybenzoate | 0.979399 |
| EX_C00954__dra | Indole-3-acetate | 0.989051 |
| EX_C00027__dra | Hydrogen peroxide | 1.130502 |
| EX_C00001__dra | H2O | 10.335535 |
| EX_C00014__dra | Ammonia | 14.375825 |
| EX_C00011__dra | CO2 | 23.827339 |

Table S5. Summary of a pFBA applied to the virgin phellogen model in heterotrophic conditions. The maximum uptake of amino acids and sucrose was fixed to 1 $mmol\cdot g_{DW}\cdot h^{-1}$, while maximizing the biomass production

| Reaction | Metabolite | Flux |
| --- | --- | --- |
| EX_C00007__dra | Oxygen | -18.443895 |
| EX_C00049__dra | L-Aspartate | -1.0 |
| EX_C00064__dra | L-Glutamine | -1.0 |
| EX_C00082__dra | L-Tyrosine | -1.0 |
| EX_C00079__dra | L-Phenylalanine | -1.0 |
| EX_C00188__dra | L-Threonine | -1.0 |
| EX_C00025__dra | L-Glutamate | -1.0 |
| EX_C00152__dra | L-Asparagine | -1.0 |
| EX_C00037__dra | Glycine | -1.0 |
| EX_C00041__dra | L-Alanine | -1.0 |
| EX_C00047__dra | L-Lysine | -1.0 |
| EX_C00089__dra | Sucrose | -1.0 |
| EX_C00065__dra | L-Serine | -1.0 |
| EX_C00062__dra | L-Arginine | -1.0 |
| EX_C00078__dra | L-Tryptophan | -1.0 |
| EX_C00123__dra | L-Leucine | -0.071343 |
| EX_C00009__dra | Orthophosphate | -0.05892 |
| EX_C00183__dra | L-Valine | -0.047559 |
| EX_C00407__dra | L-Isoleucine | -0.038312 |
| EX_C00135__dra | L-Histidine | -0.019601 |
| EX_C00073__dra | L-Methionine | -0.018207 |
| EX_C00059__dra | Sulfate | -0.014698 |
| EX_C14818__dra | Fe2+ | -2e-06 |
| EX_C00237__dra | CO | 3e-06 |
| EX_C00080__dra | H+ | 0.009598 |
| EX_Biomass__cyto | e-Biomass | 0.76682 |
| EX_C00086__dra | Urea | 0.954363 |
| EX_C00322__dra | 2-Oxoadipate | 0.95798 |
| EX_C00628__dra | 2,5-Dihydroxybenzoate | 0.979176 |
| EX_C00954__dra | Indole-3-acetate | 0.988933 |
| EX_C00027__dra | Hydrogen peroxide | 1.165448 |
| EX_C00001__dra | H2O | 9.035153 |
| EX_C00014__dra | Ammonia | 16.283406 |
| EX_C00011__dra | CO2 | 25.265562 |

Table S6. Summary of a pFBA applied to the reproduction phellogen model in heterotrophic conditions. The maximum uptake of amino acids and sucrose was fixed to 1 $mmol\cdot g_{DW}\cdot h^{-1}$, while maximizing the biomass production

| Reaction | Metabolite | Flux |
| --- | --- | --- |
| EX_C00007__dra | Oxygen | -16.711712 |
| EX_C00049__dra | L-Aspartate | -1.0 |
| EX_C00062__dra | L-Arginine | -1.0 |
| EX_C00082__dra | L-Tyrosine | -1.0 |
| EX_C00078__dra | L-Tryptophan | -1.0 |
| EX_C00025__dra | L-Glutamate | -1.0 |
| EX_C00152__dra | L-Asparagine | -1.0 |
| EX_C00037__dra | Glycine | -1.0 |
| EX_C00041__dra | L-Alanine | -1.0 |
| EX_C00065__dra | L-Serine | -1.0 |
| EX_C00089__dra | Sucrose | -1.0 |
| EX_C00064__dra | L-Glutamine | -1.0 |
| EX_C00079__dra | L-Phenylalanine | -0.98441 |
| EX_C00123__dra | L-Leucine | -0.064953 |
| EX_C00009__dra | Orthophosphate | -0.053643 |
| EX_C00183__dra | L-Valine | -0.043299 |
| EX_C00047__dra | L-Lysine | -0.038256 |
| EX_C00188__dra | L-Threonine | -0.036011 |
| EX_C00407__dra | L-Isoleucine | -0.03488 |
| EX_C00135__dra | L-Histidine | -0.017845 |
| EX_C00073__dra | L-Methionine | -0.016577 |
| EX_C00059__dra | Sulfate | -0.013381 |
| EX_C14818__dra | Fe2+ | -2e-06 |
| EX_C00237__dra | CO | 3e-06 |
| EX_C00080__dra | H+ | 0.008738 |
| EX_Biomass__cyto | e-Biomass | 0.698134 |
| EX_C00086__dra | Urea | 0.958451 |
| EX_C00628__dra | 2,5-Dihydroxybenzoate | 0.981041 |
| EX_C00954__dra | Indole-3-acetate | 0.989924 |
| EX_C00027__dra | Hydrogen peroxide | 1.240201 |
| EX_C00001__dra | H2O | 8.137847 |
| EX_C00014__dra | Ammonia | 13.444526 |
| EX_C00011__dra | CO2 | 24.290978 |
